# Editing the conserved *IPA1*–*TB1* regulatory module reshapes plant architecture and enhances tillering in wheat

**DOI:** 10.64898/2026.04.09.717480

**Authors:** Ritika Vishnoi, Gaurav Augustine, Parul Sirohi, Suchi Baliyan, Promila Choudhary, Aman Kumar, Om Prakash Raigar, Hitashi Aggarwal, Asif Islam, Jasneet Singh, Neha Kumari, Parveen Chhuneja, Ajay Kumar Pandey, Harsh Chauhan, Nitika Sandhu

**Author notes:** Ritika Vishnoi and Gaurav Augustine contributed equally and are considered as first authors. Corresponding author mail.

## Abstract

Plant architecture is a major determinant of yield potential in cereal crops, where tiller number directly influences spike production and grain yield. The transcriptional regulators Ideal Plant Architecture 1 (*IPA1*) and *Teosinte Branched1 (TB1)* constitute a conserved genetic module controlling axillary bud activity and branching in grasses; however, their functional contribution to wheat architecture remains largely unexplored. Here, we employed CRISPR/Cas9-mediated genome editing to investigate the roles of *TaIPA1* and *TaTB1* in regulating tillering in hexaploid wheat (*Triticum aestivum* L.). Comparative genomic analysis identified conserved *IPA1* orthologs across the wheat A, B, and D sub-genomes, with strong conservation of the SQUAMOSA-binding protein domain. Sequencing analysis confirmed targeted mutations, including nucleotide substitutions and insertions that generated frameshift mutations and premature stop codons. Genome-edited lines exhibited enhanced tillering compared with wild-type plants. Several *TaIPA1* mutant lines produced up to two-fold higher tiller numbers, while *TaTB1* knockout lines showed earlier tiller initiation and ∼50% increased tillering. Notably, enhanced tillering was associated with increased grain weight without affecting spikelet number per spike. Together, these results demonstrate that the conserved *TaIPA1–TaTB1* regulatory module plays a pivotal role in shaping wheat plant architecture. Targeted manipulation of this pathway using CRISPR/Cas9 provides a promising strategy for optimizing tillering and developing high-yielding wheat ideotypes.

## Introduction

Wheat (*Triticum aestivum* L.) is the cornerstone of global agriculture, contributing significantly to the world’s caloric intake and food security. As the global population continues to rise, the demand for wheat is projected to increase by 60% by 2050 (Shiferaw et al. 2013, FAO 2017). However, wheat production is facing substantial challenges from biotic and abiotic stresses, as well as the limitations of arable land (Ray et al. 2013). To meet future food demands, it is essential to enhance wheat yield potential through innovative breeding strategies.

One critical factor influencing wheat yield is tillering, the process by which the plant produces lateral shoots or tillers. Tillering directly impacts the number of spikes per plant, a key determinant of grain yield (Li et al. 2003). Optimal tillering balances the number of productive tillers while minimizing competition for resources among shoots (Mohapatra et al. 2025). However, the genetic regulation of tillering in wheat is complex and influenced by both genetic and environmental factors, necessitating targeted approaches for yield improvement.

Efforts to improve tillering in wheat have spanned traditional breeding, molecular genetics, and, more recently, genome editing. Breeding programs have historically focused on selecting high-tillering genotypes to increase the number of productive spikes per plant, thereby enhancing overall grain yield (Awan et al. 2022). The identification of quantitative trait loci (QTLs) associated with tiller number has facilitated marker-assisted selection (MAS) for optimal tillering traits, accelerating the development of superior wheat varieties (Peng et al. 2003). For instance, editing genes involved in strigolactone biosynthesis and signalling pathways has shown potential for enhancing tiller number and improving yield stability under diverse environmental conditions (Wang et al. 2021).

In recent years, advances in plant genomics have identified key regulatory genes involved in tillering. Among these, the *Ideal Plant Architecture 1* (*IPA1*) gene has garnered significant attention. Initially characterized in rice (*Oryza sativa*), *IPA1* encodes a transcription factor, *OsSPL14* that plays a pivotal role in controlling shoot architecture by regulating the expression of genes associated with tiller development and plant height (Jiao et al. 2010). Strigolactones (SLs) hormones, regulate branching inhibition through the action of *D53*, which not only suppresses the SL pathway but also directly inhibits the rice architecture regulator *IPA1* to regulate tiller number and SL-induced gene expression (Song et al. 2017). Moreover, *IPA1* directly binds to *DEP1* and *OsTB1* motifs and controlling various biological pathway including apoptosis, cell cycle, development, stress response and plant architecture in rice (Lu et al. 2013). Kerr et al. (2017) represent the SLs regulation model for shoot branching or tillering, *IPA1* controls the expression of *TB1* to inhibit tillering in rice through unknown pathways and miR156 integrates environmental/endogenous signals to post-transcriptionally suppress *IPA1* (Fig. 1a). Mutations in the *IPA1* gene have been shown to balance tiller number and plant height, leading to improved yield traits in rice (Wang and Li 2011). The functional conservation of *IPA1* across monocots suggests that its manipulation could similarly benefit wheat breeding programs (Wang et al. 2018).

**Fig. 1.**
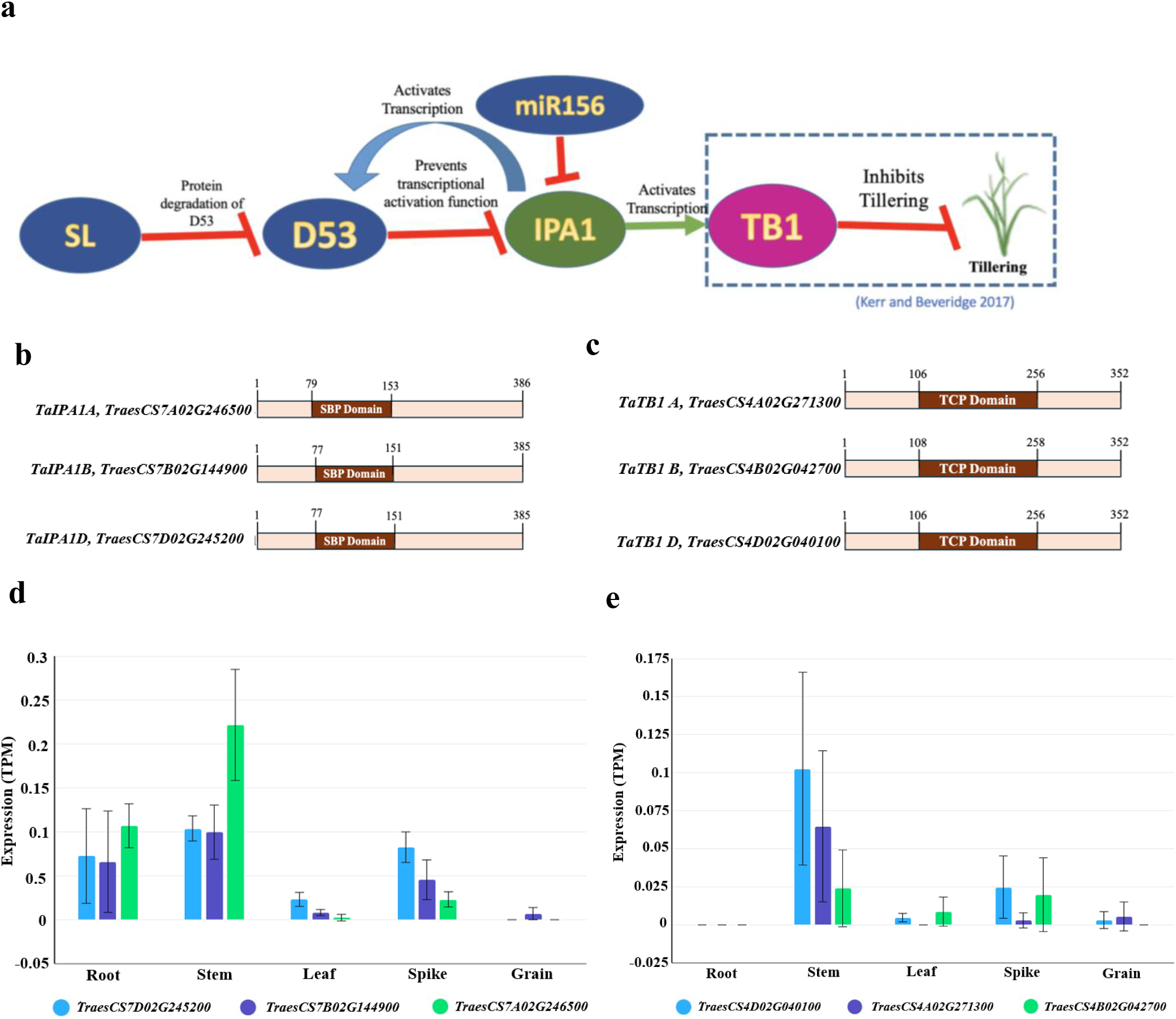
**a** *IPA1-TB1* module in tillering regulation. *IPA1*, stabilized by SL signaling via *D53/D14/D3* (preventing miR156 maturation), directly activates *TB1* transcription to repress tiller outgrowth; *TB1* knockout relieves this inhibition, enhancing tillering. **b** Schematic representation of the TaIPA1 protein domain organization in wheat. The diagram shows the conserved SBP (SQUAMOSA Promoter Binding Protein) domain, characteristic of SPL transcription factors, which is responsible for DNA binding and the regulation of downstream target genes involved in plant architecture and developmental processes. **c** Schematic representation of the *TaTB1* protein domain organization in wheat. The diagram illustrates the conserved TCP (TEOSINTE BRANCHED1/CYCLOIDEA/PCF) domain, which is characteristic of TCP transcription factors and plays a key role in DNA binding and regulation of genes controlling tillering and plant architecture. **d-e** Expression analysis of *TaIPA1* and *TaTB1* in different plant tissues, including grain, leaf, root, spike, and stem. The expression data obtained from the (http://wheatomics.sdau.edu.cn/expression/wheat.html) **d** *IPA1* **e** *TB1*

On the other hand, *Teosinte Branched1 (TB1*), a *TCP (TEOSINTE BRANCHED1/CYCLOIDEA/PCF*) family transcription is a key regulator of shoot branching and apical dominance in dicots as well as monocots. *TB1* and its orthologs play crucial roles in determining whether lateral buds develop into branches or remain suppressed, making it a focal point in studies of domestication, developmental genetics, and molecular breeding (Dixon et a. 2018). The initial discovery of *TB1’*s role in maize (*Zea mays*) domestication marked a turning point in plant evolutionary biology. Derived from the wild grass teosinte (*Zea mays ssp. parviglumis*), domesticated maize underwent dramatic architectural changes, especially in the suppression of axillary tillers and lateral branches. Doust (2007) showed that increased *TB1* expression in maize leads to enhanced apical dominance, a trait not present in teosinte, thus confirming *TB1’*s contribution to the evolution of maize plant architecture. This early work positioned *TB1* as a key gene selected during crop domestication. Since then, *TB1* and its orthologs have been functionally characterized in rice, wheat, sorghum, Arabidopsis, and perennial grasses like switchgrass and sheepgrass, highlighting a conserved regulatory framework with species-specific functional diversification influencing tillering, inflorescence patterning, and adaptation to environmental stress. *TB1* operates within a complex regulatory network that integrates hormonal signals (e.g., auxins, strigolactones, cytokinins), sugar availability, and environmental inputs such as light and nutrient status (Dong et al. 2021).

Genome editing technologies, particularly the CRISPR/Cas9 system, have revolutionized the field of plant genetics by enabling precise modifications of target genes (Jinek et al. 2012). CRISPR/Cas9 has been successfully applied to modify genes controlling agronomic traits in various crops, offering a powerful tool for enhancing crop performance (Zhang et al. 2014). For instance, *TB1* knockout in sheepgrass led to dramatic increases in tillering and biomass (Lin et al. 2023), while in-frame mutations of *OsTB1* in rice improved grain yield under phosphorus-deficient conditions (Ishizaki et al. 2023), showing that *TB1* modulation can support yield stability under nutrient stress. In wheat, the application of CRISPR/Cas9 to edit the *IPA1* and *TB1* gene presents a promising strategy to optimize tillering, thereby improving yield potential and adaptability to diverse environments.

The present work focuses on the potential of genome editing to improve wheat tillering by targeting the *TaIPA1* and *TaTB1* genes. We explore the role of *TaIPA1* and *TaTB1* in regulating shoot architecture, discuss the molecular mechanisms underlying its function, and explore the application of CRISPR/Cas9 technology in editing these genes. Furthermore, we evaluate the potential agronomic benefits of this approach and its implications for wheat breeding programs aimed at enhancing yield stability under varying environmental conditions.

## Materials and Methods

### Plant Material

The wheat cultivars C306, Unnat PBW 550 and Fielder were used for the knockout of the *TaIPA1* gene and the cultivar Fielder was used for the knockout of the *TaTB1* gene aimed at improving tillering. Unnat PBW 550 is known for its low tillering capacity under field conditions, making them a suitable candidate for enhancing this trait through genetic modification. All cultivars were cultivated under controlled conditions in a growth chamber to ensure uniform growth and minimize environmental variability.

### Expression Analysis of Candidate Genes

The expression patterns of the *TaIPA1 and TaTB1* homeologs were analysed using online tools such as WheatExp (http://wheatomics.sdau.edu.cn/expression/wheat.html) Wheat Omics, Wheat Expression, and Ensembl Plants. Spatial-temporal analyses of the homeologs were performed by retrieving data using the transcription IDs of the orthologs from the Ensembl Plants database.

### Sequencing of *TaIPA1* and *TaTB1* Gene

The coding sequence of the *TaIPA1* gene was initially extracted from the rice genome using the Ensembl database. The retrieved sequence was then aligned with the *Triticum aestivum* reference genome from the Ensembl database through BLAST analysis. Orthologous genes of *TaIPA1* were identified in the wheat reference genome (AABBDD). Three candidate *TaIPA1* genes were selected, all located on chromosome 7 of the wheat reference genome.

Specific primers were designed from the conserved regions of the three candidate genes (*TraesCS7A02G246500*, *TraesCS7B02G144900*, and *TraesCS7D02G245100*) using Primer 3 software to amplify the entire gene (Supplementary table 1). Gene amplification was carried out using NEB’s LongAmp® Taq DNA Polymerase (M0323) kit in a thermal cycler with the following program: 94°C for 30 seconds, followed by 30 cycles of 94°C for 30 seconds, 56°C for 30 seconds, and 65°C for 5 minutes, and a final extension at 65°C for 10 minutes. The amplified products were run on a 0.8% agarose gel and purified using the QIAquick PCR Purification Kit. The purified products were then sequenced using Oxford Nanopore Technology. The wheat *TaTB1-A* (*TraesCS4A02G271300*), *TaTB1-B* (*TraesCS4B02G042700*), and *TaTB1-D* (*TraesCS4D02G040100*), locus has already been sequenced (Dixon et al. 2020).

### gRNA designing for *TaIPA1* and *TaTB1* gene

Multiple sequence alignment of cDNA of the three wheat *TaIPA1* and *TaTB1* homeolog was performed using CLUSTAL Omega to identify the conserved regions among the copies. The gRNAs were designed from conserved regions shared across the homeologs to enable simultaneous targeting using WheatCrispr tool (Cram et al. 2019). Preference was given to target sites located proximal to the translational start site to increase the likelihood of generating functional knockouts. Candidate gRNAs were further evaluated for specificity by performing sequence similarity searches against the wheat reference genome using *BLAST.* Off-target potential was assessed by identifying genomic regions with partial sequence similarity to the gRNA. The selected gRNAs exhibited minimal predicted off-target activity, with no genomic sites showing high-stringency matches beyond short stretches of sequence similarity. Based on these criteria, two high-specificity gRNAs were selected for the development of *TaIPA1* and *TaTB1* knockout lines.

### Construction of CRISPR-Cas9 vector with sgRNA

The JD633 Cas9 vector with *GRF4-GIF1* (Debernardi et al. 2020; Tyagi et al., 2026) was used for editing in hexaploid wheat. For preparing the cassette, JD633 vector was digested with *Aar*1 restriction enzyme at 37 °C for 3 hours and removed 456bp fragment from vector. sgRNA oligo duplex prepared using reaction mixture, forward oligo 4μl, reverse oligo 4 μl, Phusion HF buffer 2 μl in thermal cycler with following programme 37 °C for 60 min, 95 °C for 10 min and Cool down @ 0.1 °C to 25°C. 100 times diluted product was used for ligation reaction with digested JD633 plasmid by T4 DNA ligase kit (NEB) at 16°C for 5 hours. 5μl of ligated product was used for transformation of competent DH5α cells by heat shock method at 42°C. Further, construction of CRISPR/Cas9 vector with sgRNA was confirmed by Sanger Sequencing. Finally, confirmed plasmid was transformed in *AGL1* Agrobacterium strain for the transformation of wheat.

### *Agrobacterium*-mediated transformation in wheat cultivars

*Agrobacterium*-mediated wheat transformation was performed following the protocol of Hayta et al. (2021) with minor modifications, using immature embryos. Immature seeds were sterilized with 70% ethanol for 2 min followed by 4% sodium hypochlorite for 10 min and rinsed five times with sterile distilled water. Immature embryo was isolated and infected with an *Agrobacterium* suspension for 20 min followed by co-cultivation on medium at 24 ± 1°C in the dark for 5 days. After co-cultivation, the embryogenic axes were excised and cultured on wheat callus induction (WCI) medium supplemented with 160 mg/liter Timentin for 5 days. Hygromycin selection (15 mg L⁻¹) was applied for two cycles of two weeks each. The surviving calli were transferred to regeneration medium containing 2.0 mg/liter Zeatin and incubated under light (16h) and dark condition (8h) for 14 days to induce shoot regeneration. Regenerated plantlets were transferred to MS rooting medium and subsequently acclimatized in soil pots maintained at 70–80% relative humidity.

### PCR and Sequencing-Based Identification of Mutants

Genomic DNA was isolated from the putative transgenic wheat plants using the standard CTAB extraction method. The quality and integrity of the DNA were assessed by agarose gel electrophoresis, and the concentration was quantified using a nanodrop spectrophotometer. For molecular screening, PCR amplification was carried out using primers specific to the selectable marker gene *hptII*. Plants producing the expected *hptII* amplicon were considered putative transformants and were subsequently selected for further molecular analysis. To identify the mutations at target region of *TaIPA1* and *TaTB1* genes, the genome specific primers were used for the PCR amplification. The target region of *TaIPA1* and *TaTB1* genes were amplified from the *hptII*-positive lines. The amplified PCR fragments were purified to remove excess primers and dNTPs, and subsequently subjected to Sanger sequencing. Sequencing results were analysed to confirm the successful editing events at the target regions of the *TaIPA1* and *TaTB1* genes in the transformed plants. Furthermore, to examine the homeolog specific editing, primers were designed from the conserved region of the three copies of *TaIPA1* and *TaTB1* genes and cloned in *ECoR*V linearised pbS SK(+) vector. Several colonies were sequenced to identify the editing events across A, B and D genomes.

## Results

### In *silico* identification and sequence characterization of *TaIPA1* and *TaTB1* orthologs in hexaploid wheat

To identify orthologs of the rice *IPA1* gene in hexaploid wheat, the rice *OsIPA1* sequence was retrieved and used as a query for BLAST analysis against ten wheat genome assemblies available on the Ensembl Plants database. The search identified highly similar sequences located on wheat chromosomes 5A, 5B, 5D, 7A, 7B, and 7D. Among these, the closest wheat orthologs were *TraesCS7A02G246500*, *TraesCS7B02G144900*, and *TraesCS7D02G245200*, which showed approximately 50% sequence similarity with rice *OsIPA1* and ∼80% similarity among the wheat orthologs, indicating strong conservation across the wheat sub-genomes (Supplementary Fig. 1a).

Domain comparison between the homoeologous wheat transcripts (7A, 7B.1, 7B.2, 7D.1, and 7D.2) and rice *OsIPA1* revealed conservation of the SQUAMOSA-binding protein (SBP) domain (Fig. 1b). Protein sequence alignment further confirmed the conservation of this domain, suggesting that a single gRNA could potentially target all three homeologs (Supplementary Fig. 1b). Phylogenetic analysis clustered the sequences according to their respective sub-genomes, indicating their shared evolutionary origin and functional conservation.

The wheat *TaTB1* orthologs corresponding to the A, B and D sub-genomes have previously been characterized, including *TaTB1-A* (*TraesCS4A02G271300*), *TaTB1-B* (*TraesCS4B02G042700*), and *TaTB1-D* (*TraesCS4D02G040100*) (Dixon et al., 2018). These genes encode TCP transcription factors known to regulate tillering and spike architecture in wheat (Fig. 1c). The availability of well-annotated sequences for all three *TaTB1* homeologs facilitated the design of guide RNAs targeting conserved regions across the sub-genomes for genome editing. As the sequences of the three *TaTB1* homeologs are already well annotated in the wheat reference genome, sequencing efforts in this study focused on the *TaIPA1* orthologs in the wheat cultivar Unnat PBW550. Genome-specific primers targeting the *TaIPA1* loci on chromosomes 7A, 7B, and 7D were designed based on the IWGSC RefSeq v2.0. Genomic fragments (500 bp to 1.5 kb) were amplified and sequenced from Unnat PBW550 via Nanopore sequencing. The resulting sequences were deposited in the NCBI GenBank (PP069455, PP069456, PP069457, PP069458, PP069459, PP069460) (Supplementary Fig. 2). Sequence alignment showed **∼**99% similarity with the Chinese Spring reference genome, indicating strong conservation of *TaIPA1* gene, particularly exon 1, supporting its suitability as a CRISPR/Cas knockout target.

### Expression analysis of *TaIPA1* and *TaTB1* during wheat development

To understand their potential roles in wheat development and tillering regulation the expression patterns of *TaIPA1* and *TaTB1* genes were analysed during the growth and across different wheat tissues. The transcript abundance across different tissues (root, shoot, leaf, spike, and grain) was retrieved from the wheat expression database (http://wheatomics.sdau.edu.cn/expression/wheat.html) for the cultivar Chinese Spring. The results indicated that *TaIPA1* and *TaTB1* orthologs are predominantly expressed in stem and spike tissues, suggesting their involvement in spike and shoot architecture during early reproductive development (Fig. 1d-e).

### Wheat Transformation for generation of knockout mutants

To perform CRISPR based genome editing in wheat, one highly efficient gRNA for each target gene was cloned into the vector JD633 (Fig. 2 and Supplementary Fig. 3) and was used for the *Agrobacterium*-mediated transformation of wheat cultivars cv. Fielder, C306, and Unnat PBW550. A total of 24 independent putative mutants of Fielder and 4 of Unnat PBW550 were generated for *TaIPA1* gene, while 8 putative mutants in Fielder were generated for *TaTB1* gene (Supplementary Fig. 4). PCR screening using *hptII*-specific primers confirmed T-DNA integration in the transformed plants. For *TaIPA1*, 8 lines of Fielder and 1 of Unnat PBW550 were positive for the integration of T-DNA (Fig. 3a(i)). Similarly, for *TaTB1*, seven plants for fielder showed successful T-DNA insertion (Fig. 3b(i)). All the PCR-positive plants were subsequently subjected to targeted sequencing to confirm CRISPR-induced edits at the respective genomic loci.

**Fig. 2.**
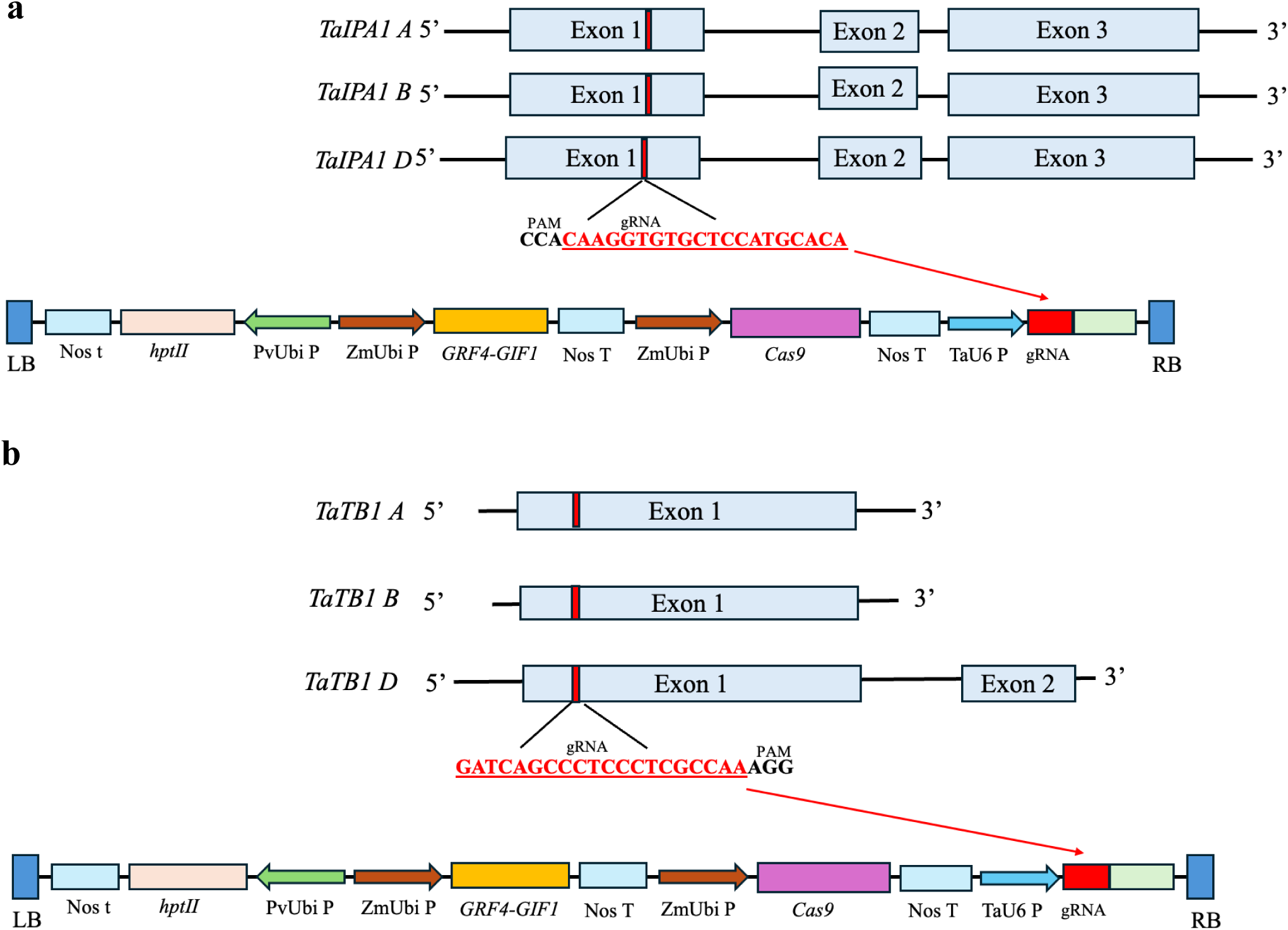
CRISPR–Cas9 guide RNA (gRNA) construct design targeting the three homeologous copies of **a** *TaIPA1* and **b** *TaTB1* genes in hexaploid wheat. The gRNAs were designed to bind conserved sequences shared among the A, B, and D subgenomes, enabling simultaneous genome editing of all homeologs. A red mark on the exon, and the underlined red sequence represents the guide RNA; the underlined black sequence represents the PAM sequence.

**Fig. 3.**
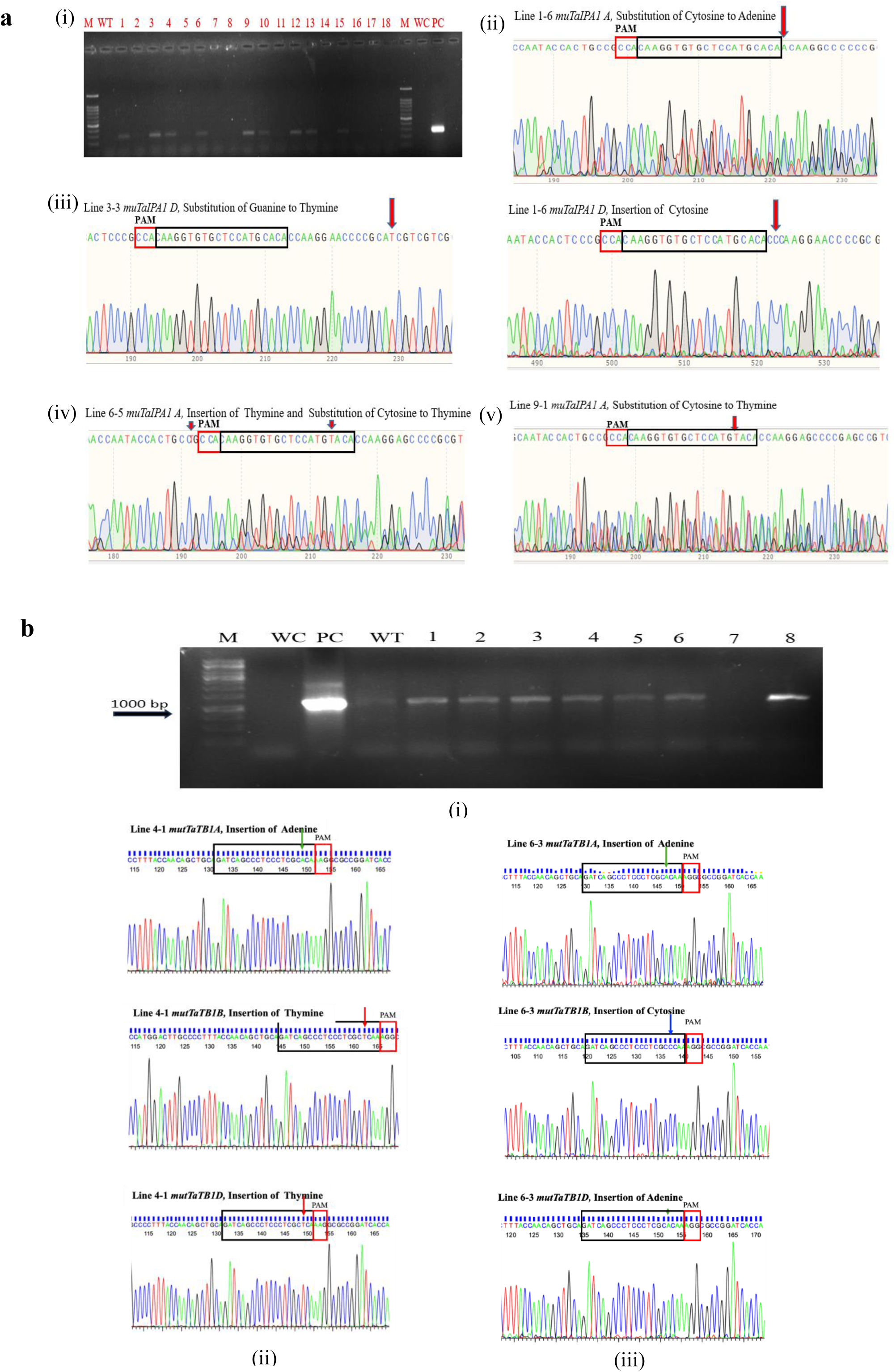
Molecular confirmation of the transgenic lines of *TaIPA1* (**a**) and TaTB*1* (**b**). **a**(i)PCR-based confirmation of putative transgenic plants using the *hygromycin phosphotransferase II (hptII)* gene. **a**(ii) Sequencing chromatogram of Target genomic region of *TaIPA1* mutant line 1-6 showing substitution of cytosine to adenine in A copy and insertion of cytosine in D copy. **a**(iii) Sequencing chromatogram of Target genomic region of *TaIPA1* mutant line 3-3 showing substitution of guanine to thymine in D copy. **a**(iv) Sequencing chromatogram of Target genomic region of *TaIPA1* mutant line 6-5 showing substitution of cytosine to thymine in A copy. **a**(v) Sequencing chromatogram of Target genomic region of *TaIPA1* mutant line 9-1 showing substitution of cytosine to thymine in A copy. **b**(i) PCR-based confirmation of putative transgenic plants using the *hygromycin phosphotransferase II (hptII)* gene. **b**(ii) Chromatogram showing insertion of adenine base in A copy of *TaTB1* and thymine in B and D copies in transgenic line 4-1. **b**(iii) Chromatogram showing insertion of adenine base in A and D copies of *TaTB1*, cytosine in B in transgenic line 6-3.

### Genotypic characterization of *TaIPA1* and *TaTB1* mutants

The *TaIPA1* target region was amplified from the genomic DNA of putative mutants using IPA1F and AR1 primers with NEB Q5 High-Fidelity DNA Polymerase, followed by digestion with T7 endonuclease. In the putative mutant plants (P2, P4, P5, and P6), additional cleavage fragments were detected along with the expected 730 bp amplicon, indicating the presence of heteroduplex mismatches generated by CRISPR/Cas9-induced mutations. In contrast, wild-type plants (Fielder and Unnat PBW550) showed only the intact 730 bp band, confirming the absence of mutations. Among the 29 screened plants, mutations were detected in four plants P2, P4, P5, P6 in the Fielder background (Supplementary Fig. 5). Seeds from the confirmed T_0_ plants were advanced to T_1_ generation and grown in containment conditions. High tillering T1 lines were further analysed by sanger sequencing to characterize edits at the target locus.

Sequence analysis revealed single-nucleotide substitutions (C→A, G→A, and C→T) as well as single-base insertions within the *TaIPA1* target region (Fig. 3a (ii–v); Supplementary Table 2). Some of these insertions resulted in frameshift mutations, and such lines were selected as putative *TaIPA1* knockdown/knockout candidates for further analysis.

For the *TaTB1* gene, sequencing of putative mutants identified single-base insertions (adenine or thymine) at the target site located four bases upstream of the PAM sequence (Supplementary Table 3; Supplementary Fig. 6). These insertions caused frameshift mutations within the coding region of *TaTB1*. These frameshift mutations introduced premature stop codons, producing truncated TB1 proteins of 168–170 amino acids compared with the full-length wild-type proteins (352–354 aa). These insertions resulted in disturbing the TCP domain of TB1 protein and hence altering its function. Seeds from edited plants were advanced to the next generation, and sequencing of two high-tillering progeny confirmed the presence of stable CRISPR/Cas9-induced edits across the three genomes and suggesting complete *TaTB1* knockout due to introduction of premature stop codon (Fig. 3b (ii–iii); Supplementary Table 4). These edited plants were therefore selected for further characterization.

### Phenotypic characterization of *TaIPA1* and *TaTB1* mutants

The tillering capacity of the mutant lines was evaluated at the T₁ stage in comparison with the wild-type control (Fielder), revealing substantial variations among the evaluated lines (Fig. 4a-c). Among the edited lines, mutTaIPA1_1-6 and mutTaIPA1_3-3 exhibited the highest tiller numbers, producing an average of 6 tillers per plant, followed by mutTaIPA1_9-1 with 5 tillers. Line mutTaIPA1_6-5 showed moderate tillering with an average of 4 tillers per plant. In contrast, the wild-type Fielder produced approximately 3 tillers per plant. The enhanced tillering observed in mutTaIPA1_1-6, mutTaIPA1_9-1, mutTaIPA1_3-3, and mutTaIPA1_6-5 indicates that disruption of the *TaIPA1* regulatory pathway may promote axillary bud outgrowth and increase tiller production.

**Fig. 4.**
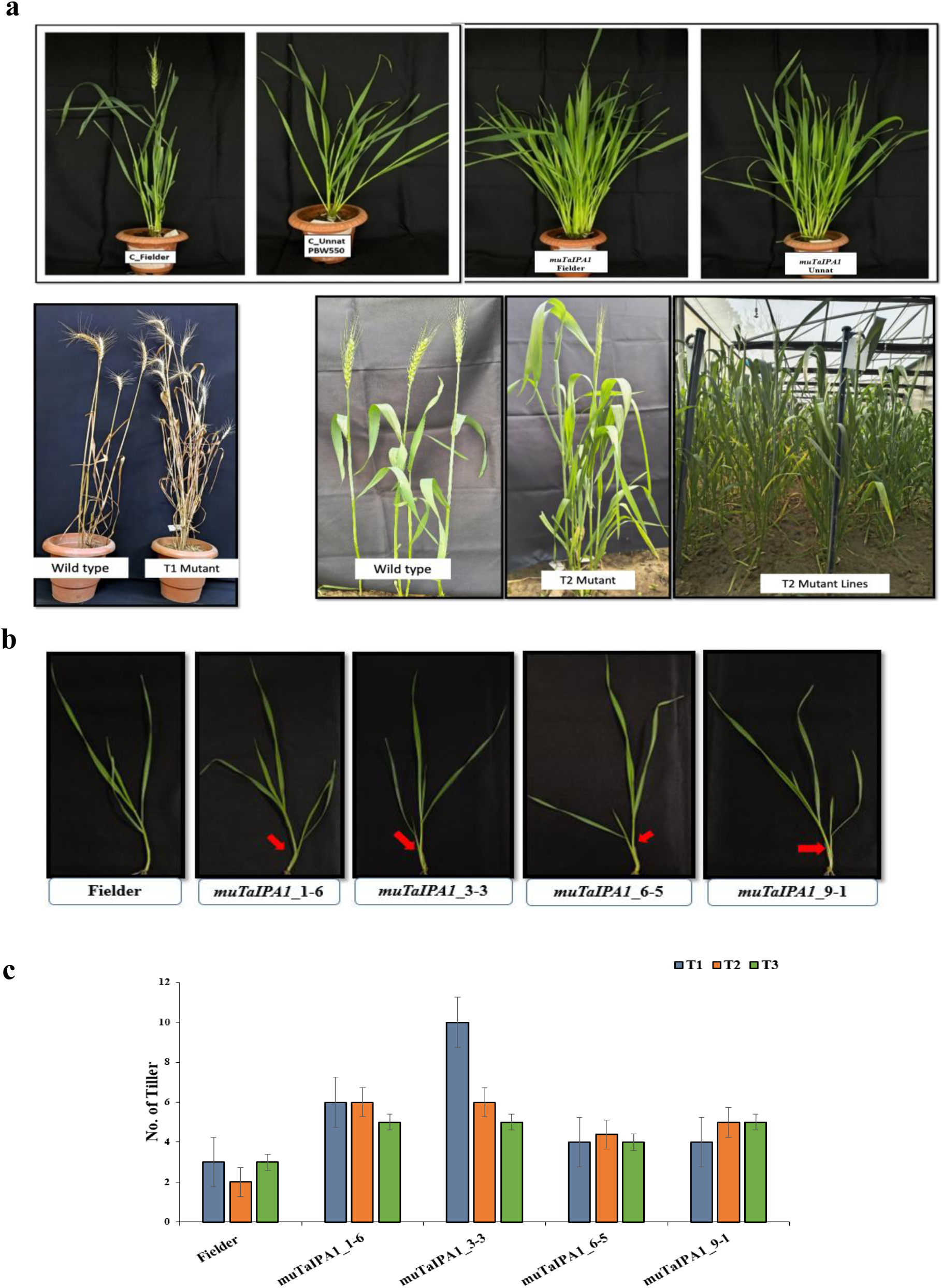
Phenotypic analysis of *TaIPA1* knockout lines. **a** Mutant lines produce more tillers than the wild type. **b** Initiation of tiller in wild type and mutant lines at early stage. **c** Number of tillers in wild type and mutant lines in T_1_, T_2_ and T_3_ generation.

To further investigate the role of *TaTB1* in regulating shoot branching, progenies derived from confirmed knockout plants were evaluated for tillering performance. A distinct phenotypic difference was observed during vegetative growth, where *TaTB1* knockout plants-initiated tiller development earlier than the wild type (Fig. 5a-b). This early initiation suggests that the loss of *TaTB1* function accelerates axillary bud activation, consistent with the well-established role of *TaTB1* as a negative regulator of tiller development and apical dominance.

**Fig. 5.**
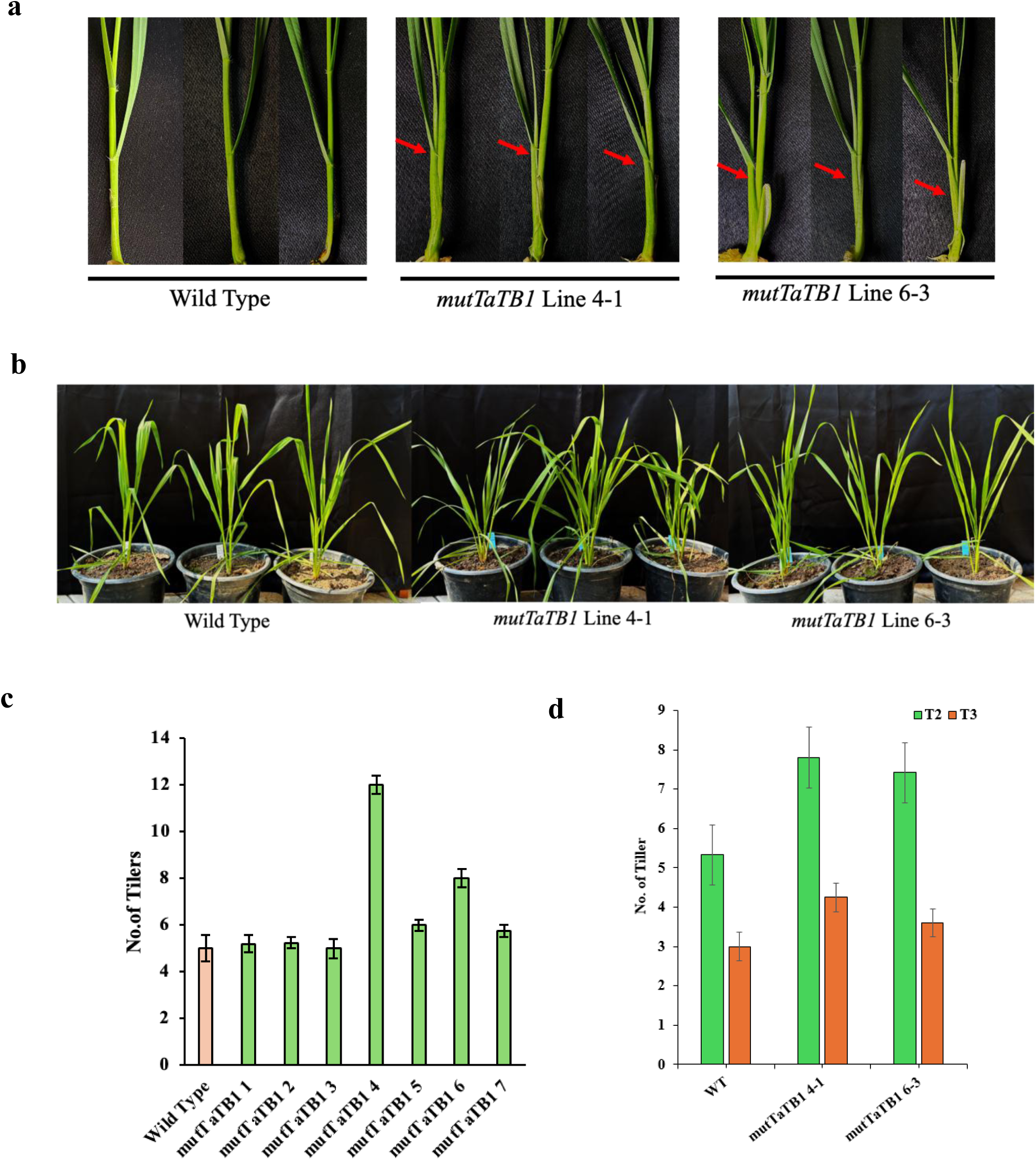
Phenotypic analysis of *TaTB1* knockout lines. **a** Mutant lines showing early onset of tiller initiation than wild-type. **b** Mutant lines produce more tillers than the wild type. Number of tillers in wild type vs *TaTB1* knockout lines **c** T_1_ **d** T_2_, and T_3_ generation. Error bar denotes the standard error, where n=8.

Among the evaluated lines, knockout line 4 exhibited the most pronounced phenotype, producing approximately 50% more tillers than the wild-type plants (Fig. 5c). These findings highlight the functional importance of *TaTB1* in regulating shoot architecture and identify line 4 as a promising genetic resource for improving tillering potential in wheat breeding programs. The agronomic implications of these mutations were further examined by assessing grain weight and the average number of spikelets per spike (Supplementary Table 5). Several mutant lines exhibited higher grain weight compared with the wild-type Fielder, suggesting a potential yield advantage. However, no significant differences were observed in spikelet number per spike among the evaluated lines, indicating that the mutations primarily influenced tillering without adversely affecting spikelet formation.

## Discussion

Optimization of plant architecture has been a major target during cereal domestication and modern crop improvement. Among architectural traits, tiller number plays a central role in crop productivity as it determines the spike number per plant which determines the yield potential in cereal crops. Tiller development is a highly coordinated biological process controlled by transcriptional regulators, hormonal signalling pathways, and environmental cues. Among these key regulators, the *TaIPA1-TaTB1* regulatory module represents a conserved pathway controlling axillary bud activity and branching potential in grasses. In the present study, CRISPR/Cas9 mediated knockout of *TaIPA1* and *TaTB1* in hexaploid wheat resulted in high-tillering phenotypes, indicating that these genes act as key regulators of wheat plant architecture.

*TaIPA1* (Ideal Plant architecture 1), also known as SPL14 in rice encodes a Squamosa Promoter Binding protein-like (SPL) transcription factor. Previous studies have demonstrated that *TaIPA1* negatively regulates tiller development by modulating the expression of genes involved in axillary bud initiation and growth (Lu et al. 2013). In cereals, *TaIPA1* integrates hormonal signals-particularly strigolactone and cytokinin to fine-tune the branching patterns. Through these interactions, *TaIPA1* maintains a balance between apical dominance and lateral branching, ensuring optimal allocation of assimilates between vegetative and reproductive structures (Li et al. 2022).

Our findings align with earlier reports showing that overexpression of *TaSPL14* reduces tiller number in wheat (Jian et al. 2024). In contrast, CRISPR/Cas9 mediated disruption of *TaIPA1* in present study using developmental regulators cassette (GRF4-GIF1 module) resulted in a marked increase in generating edited lines in diverse cultivars with enhanced tiller number relative to wild-type plants. Previously, such module were shown to overcome the genotypic bottlenecks for wheat transformation (Tyagi et al., 2026).

The high tillering observed in the mutant lines was accompanied by improved yield performance, suggesting that that the additional tillers were productive and contributed to spike formation. This observation indicated that the loss of *TaIPA1* function releases the repression on axillary bud outgrowth without imposing detrimental trade-off between wheat development. Such architectural modification is particularly relevant for improving yield potential under high-density planting systems, where optimized tillering can enhance canopy productivity. In parallel, *Teosinte Branched 1 (TB1)* represents another key transcriptional regulator of branching in grasses. *TB1* was originally identified as a major domestication gene in maize, where it played a critical role in reducing lateral branching and enhanced apical dominance (Doebley et al. 1995; Zhou et al. 2011). In wheat *TaTB1* functions as a transcriptional repressor of axillary bud outgrowth (Lewis et al. 2008). In our study, CRISPR/Cas9 generated *TaTB1* knockout line exhibited increased tiller number further confirming the conserved role of *TaTB1* in regulating plant architecture. The increased tillering phenotype observed in the *TB1* knockout lines are consistent with previous studies across cereal species. Genetic and molecular studies have established that *TB1* acts as downstream of strigolactones signalling and integrates hormonal signals to regulate bud dormancy (Dun et al. 2023). In rice, strigolactones signalling promotes *OsTB1* expression, which in turn suppresses axillary bud growth (Takeda et al. 2003; Dun et al. 2011). Similar regulatory functions of *TB1* homologs have been reported in barley, sorghum, and other grasses, underscoring the evolutionary conservation of this pathway in controlling branching architecture (Dixon et al. 2018). The results obtained in this study therefore reinforce the conserved nature of *TB1*-mediated regulation of tillering across cereal crops.

Importantly, *TaIPA1* and *TaTB1* operate within an interconnected regulatory network rather than functioning independently. Evidence from previous studies suggests that *TaIPA1* can influence *TaTB1* expression either directly or indirectly, and both the genes converge on hormonal pathways that regulate axillary bud activation. This coordinated regulation framework enables plants to precisely modulate branching pattern in response to developmental stages and environmental conditions. The *IPA1*-*TB1* module therefore acts as a key architectural switch that determine the balance between apical dominance and lateral growth in grasses. Disruption of either component of this module releases axillary buds from dormancy, resulting in increased tiller formation.

Another important implication of our study is the demonstration of efficient genome editing in hexaploid wheat genome. Because the targeted genes are highly conserved across three wheat sub genomes, a single guide RNA enabled simultaneous editing all homeologous copies. This highlights the precision and effectiveness of CRISPR/Cas9 technology for functional genomics studies and targeted trait improvement in polyploid crops. The ability to edit multiple homeologs simultaneously represents a major advantage for accelerating genetic improvement in wheat, where gene redundancy often masks phenotypic effects.

From a breeding perspective, the architectural modification observed in the edited lines provides valuable insights for designing high-yielding wheat ideotypes. Optimizing tiller number without compromising spike fertility or grain weight is a key objective in wheat breeding. The results presented here suggest that targeted manipulation of *TaIPA1* and *TaTB1* can be used to fine-tune tillering while maintaining productive spike development. Such architectural optimization could improve resource-use efficiency, light interception, and canopy structure, thereby enhancing yield potential under changing agro-ecological conditions. Furthermore, the generation of marker-free knockout lines represents an important step toward the practical deployment of genome-edited crops. The edited lines developed in this study therefore provide valuable genetic resources that could be directly utilized in breeding programs aimed at improving wheat productivity

## Conclusion

This study highlights the conserved and fundamental role of the *IPA1*-*TB1* regulatory module in controlling tillering and plant architecture in wheat. Both *TaIPA1* and *TaTB1* act as key transcriptional repressors that limit axillary bud outgrowth and maintain apical dominance by integrating hormonal signalling pathways. CRISPR/Cas9-mediated disruption of these genes resulted in enhanced tillering and improved productivity, demonstrating their strong potential as targets for architectural optimization in wheat improvement. Strategic manipulation of the *IPA1*-*TB1* pathway therefore represents a promising approach for developing high-yielding wheat varieties capable of meeting the growing global demand for food.

## DATA AVAILABILITY STATEMENT

The additional material is available in supplementary material.

## ACKNOWLEDGMENTS

The work was compiled under the projects funded by the Department of Biotechnology, Govt. of India (Grant No. BT/PR36830/GET/119/343/2020).

## AUTHOR CONTRIBUTIONS

NS, HC, AP and PC conceived the project, designed the study, and provided the necessary resources. Experimental work, including gene sequencing and vector construction, was carried out by GA, RV, AK, PCO, and JS. Wheat transformation, qRT-PCR analysis, and genotyping were performed by GA, RV, AK, HA, PS, SB, AI, and NK. Phenotyping was performed by RV, GA, and OPR. Data analysis and compilation of genotypic and phenotypic datasets were conducted by RV and GA. The manuscript was written by GA, RV, and NS. NS, HC, AP, and PC critically revised the manuscript. All authors read and approved the final version of the manuscript.

## Declarations

The authors declare that they have no conflict of interest.

---

**Supplementary Table 1.**
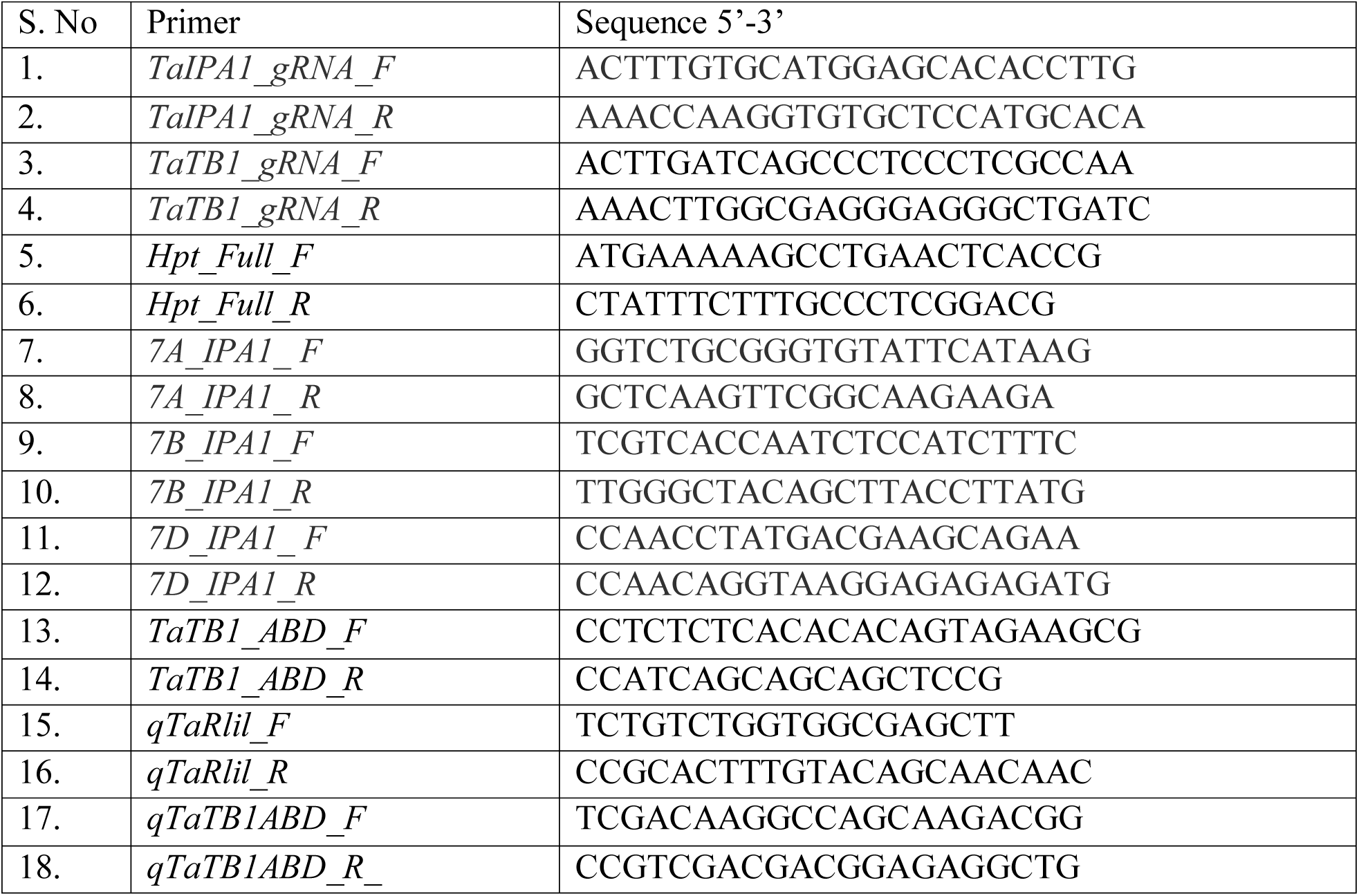
List of primers used in the study.

**Supplementary Table 2.**
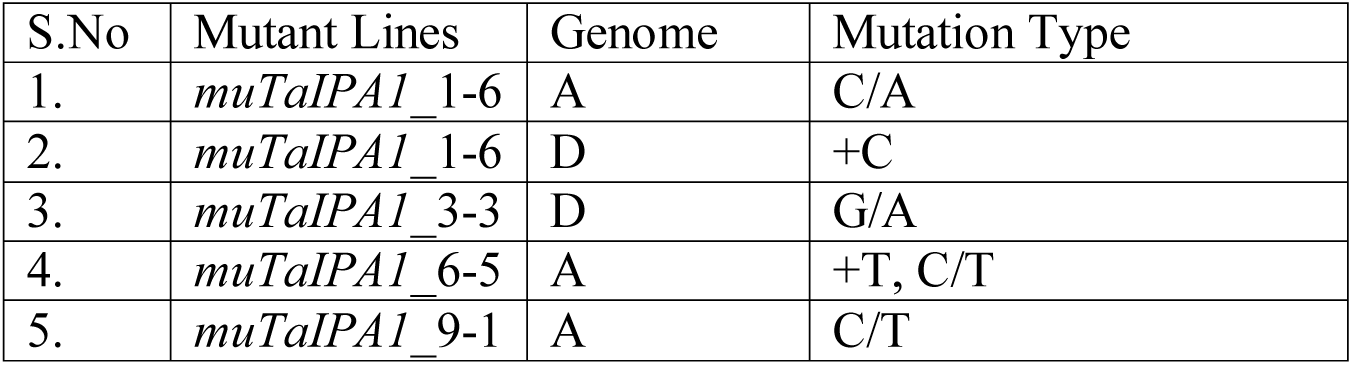
Description of Mutation in edited lines of *TaIPA1*.

**Supplementary Table 3.** Description of Mutation on the D copy of *TaTB1* gene and effect on protein in edited lines of *TaTB1*.

| S.No | Mutant Lines | Genome | Mutation Type | Effect on protein |
| --- | --- | --- | --- | --- |
| 1. | Line 1 | D | Insertion of Adenine | Premature stop codon introduced |
| 2. | Line 2 | D | Insertion of Thymine | Premature stop codon introduced |
| 3. | Line 3 | D | Insertion of Thymine | Premature stop codon introduced |
| 4. | Line 4 | D | Insertion of Thymine | Premature stop codon introduced |
| 5. | Line 5 | D | Insertion of Thymine | Premature stop codon introduced |
| 6. | Line 6 | D | Insertion of Adenine | Premature stop codon introduced |
| 7. | Line 7 | D | Insertion of Thymine | Premature stop codon introduced |

**Supplementary Table 4.** Description of Mutation on all three homeologs of *TaTB1D* and effect on protein in edited lines of *TaTB1*.

| S.No | Mutant Lines | Genome | Mutation Type | Effect on protein |
| --- | --- | --- | --- | --- |
| 1. | Line 4-1 | A | Insertion of Adenine | Premature stop codon introduced |
| 2. | Line 4-1 | B | Insertion of Thymine | Premature stop codon introduced |
| 3. | Line 4-1 | D | Insertion of Thymine | Premature stop codon introduced |
| 4. | Line 6-3 | A | Insertion of Adenine | Premature stop codon introduced |
| 5. | Line 6-3 | B | Insertion of Cytosine | Premature stop codon introduced |
| 6. | Line 6-3 | D | Insertion of Adenine | Premature stop codon introduced |

**Supplementary Table 5.**
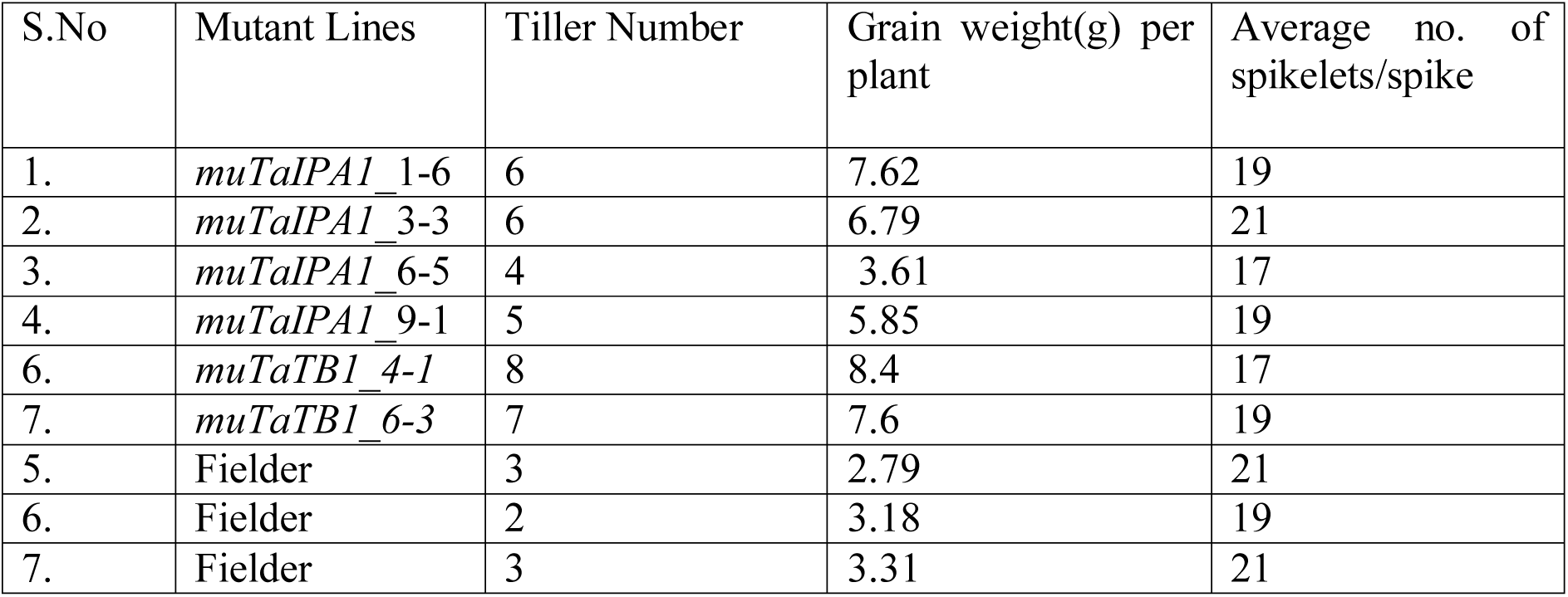
Tiller number, grain weight, Average no. of spikelets/ spike in the genome edited mutant lines for *TaIPA1* at T_2_ generation.

**Supplementary Figure 1.**
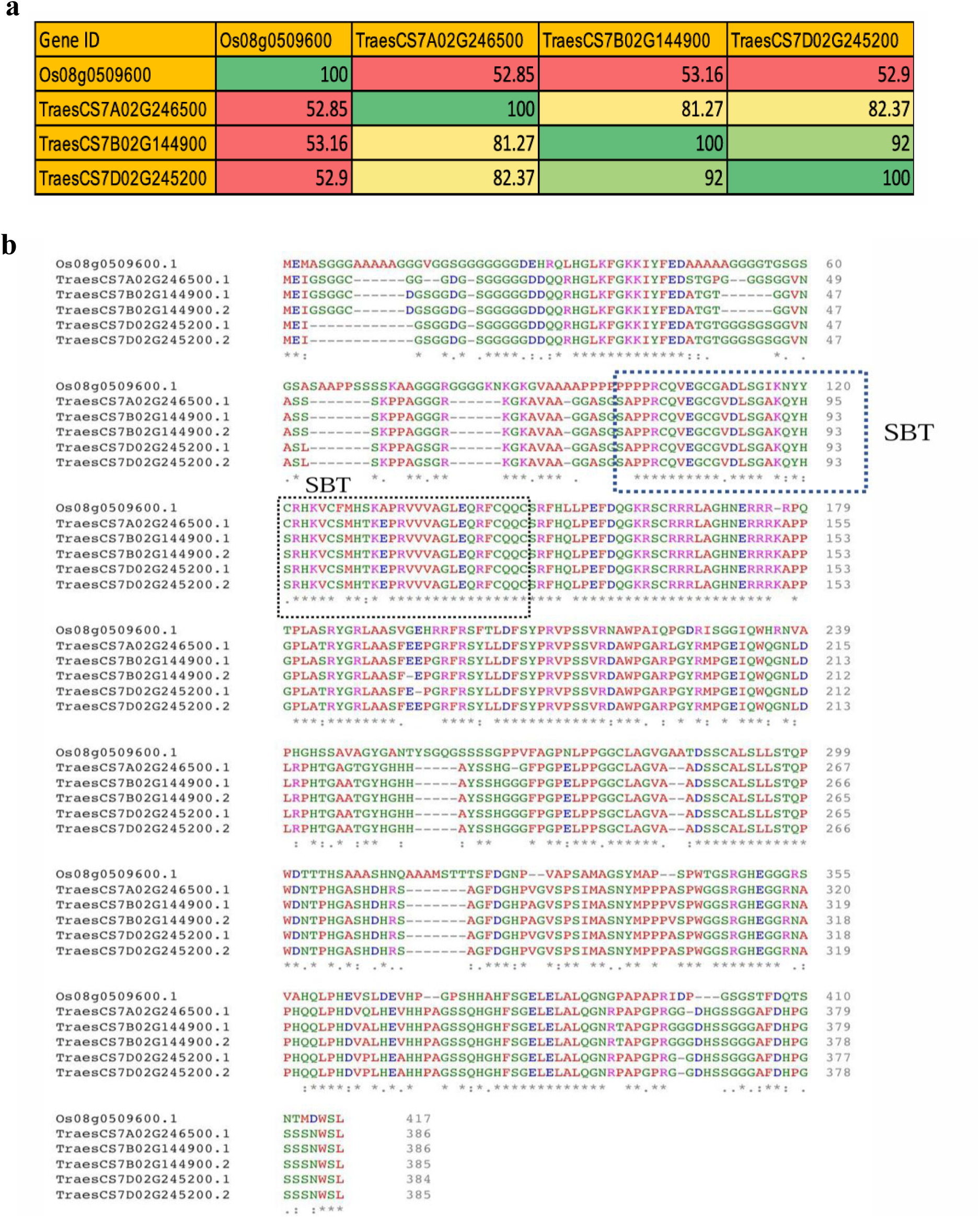
**a** Percentage identity matrix for multiple sequence alignment of rice IPA1 (*Os08g0509600*) and the wheat orthologs (*TraesCS7A02G246500, TraesCS7B02G144900 and TraesCS7D02G245200*). Color codes representing the highest identity percentage with green color, while the lowest with red color. **b** Protein sequence alignment for the rice IPA1 protein sequence (*Os08g0509600.1*) and the three wheat IPA1 homoeolog protein sequences and their transcripts. The area in the boxes denotes the conserved SBT domain

**Supplementary Figure 2.**
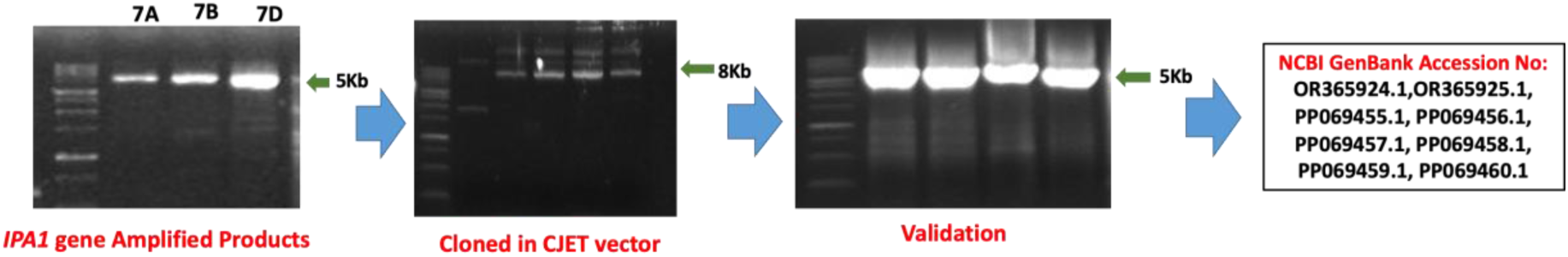
Cloning of *TaIPA1* in CloneJet, validation and submission in NCBI database.

**Supplementary Figure 3.**
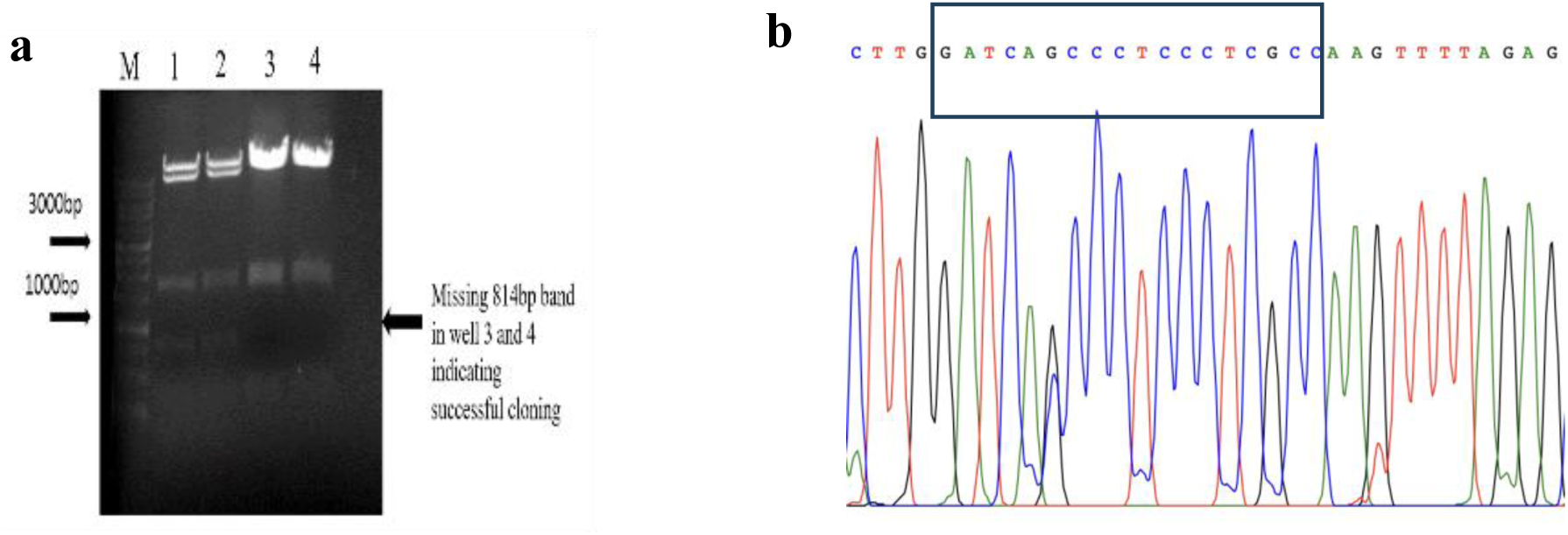
Confirmation of gRNA cloning. **a** Agarose Gel electrophoresis validating the cloning of gRNA in JD633. **b** Chromatogram showing the gRNA in JD633.

**Supplementary Figure 4.**
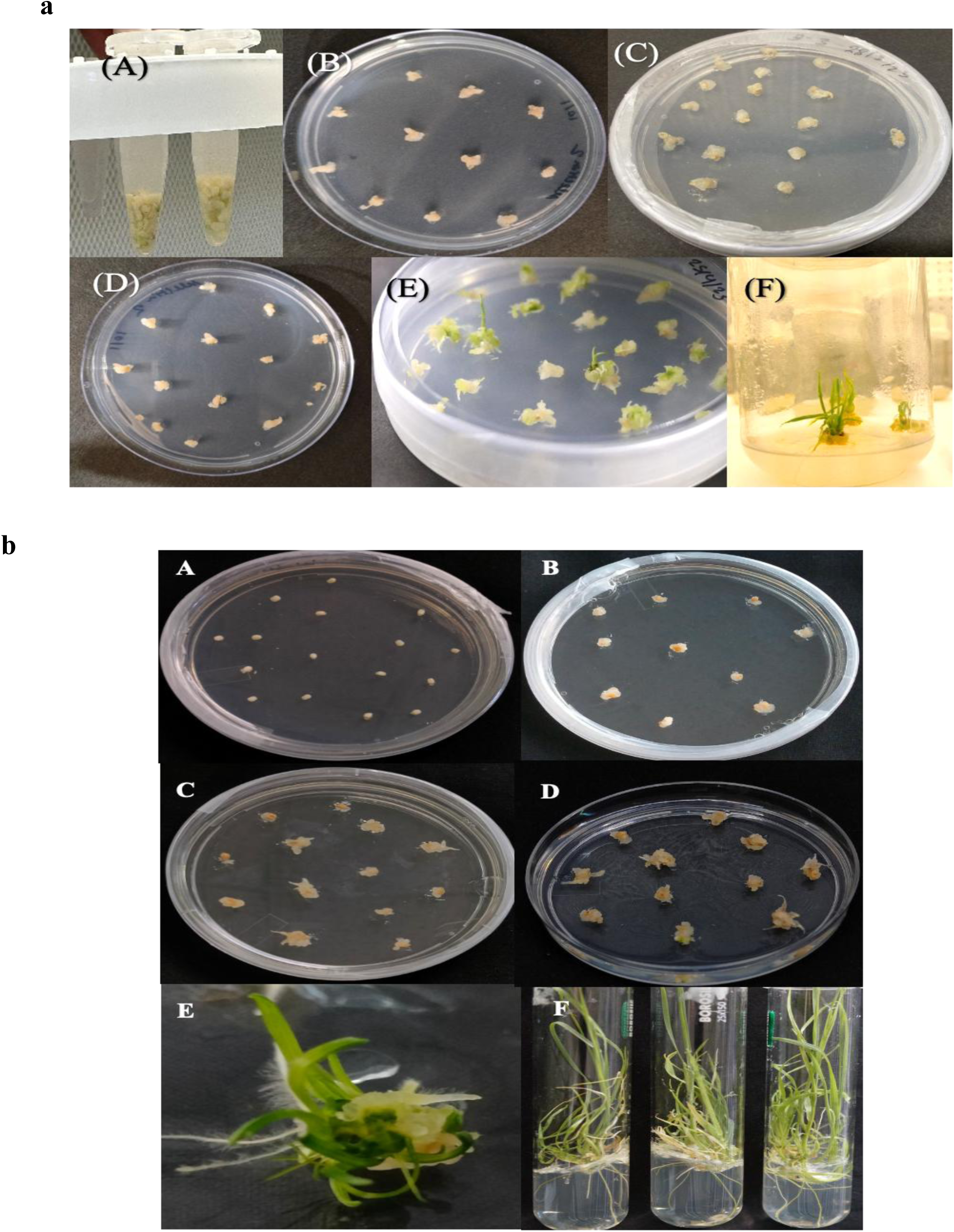
Agrobacterium-mediated transformation of wheat immature embryos for the development of knockout lines of (**a**) *IPA1* (**b**) *TaTB1*.

**Supplementary Figure 5.**
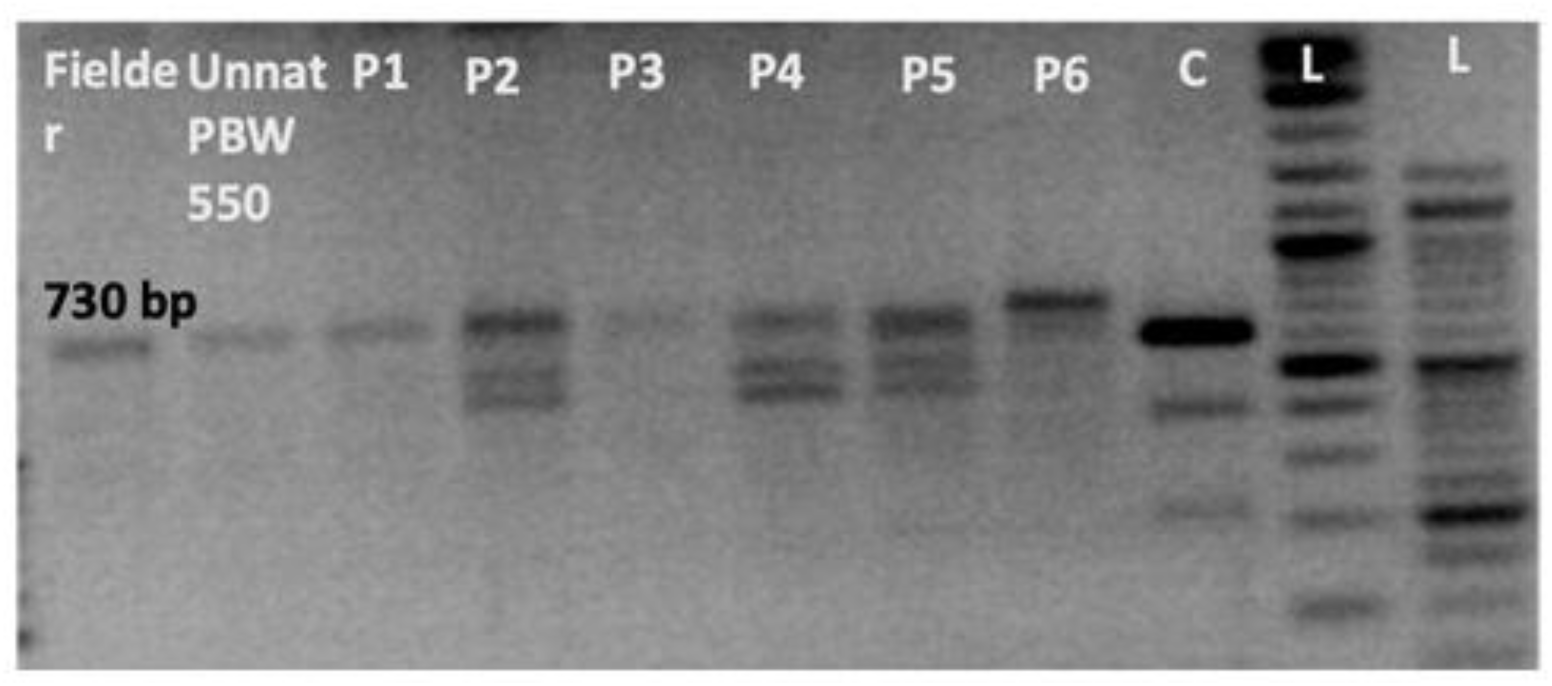
Detection of mutations in the *TaIPA1* gene using the T7 endonuclease I assay. Fielder and Unnat PBW550 were used as wild-type controls, while P1–P6 represent putative mutant plants.

**Supplementary Figure 6.**
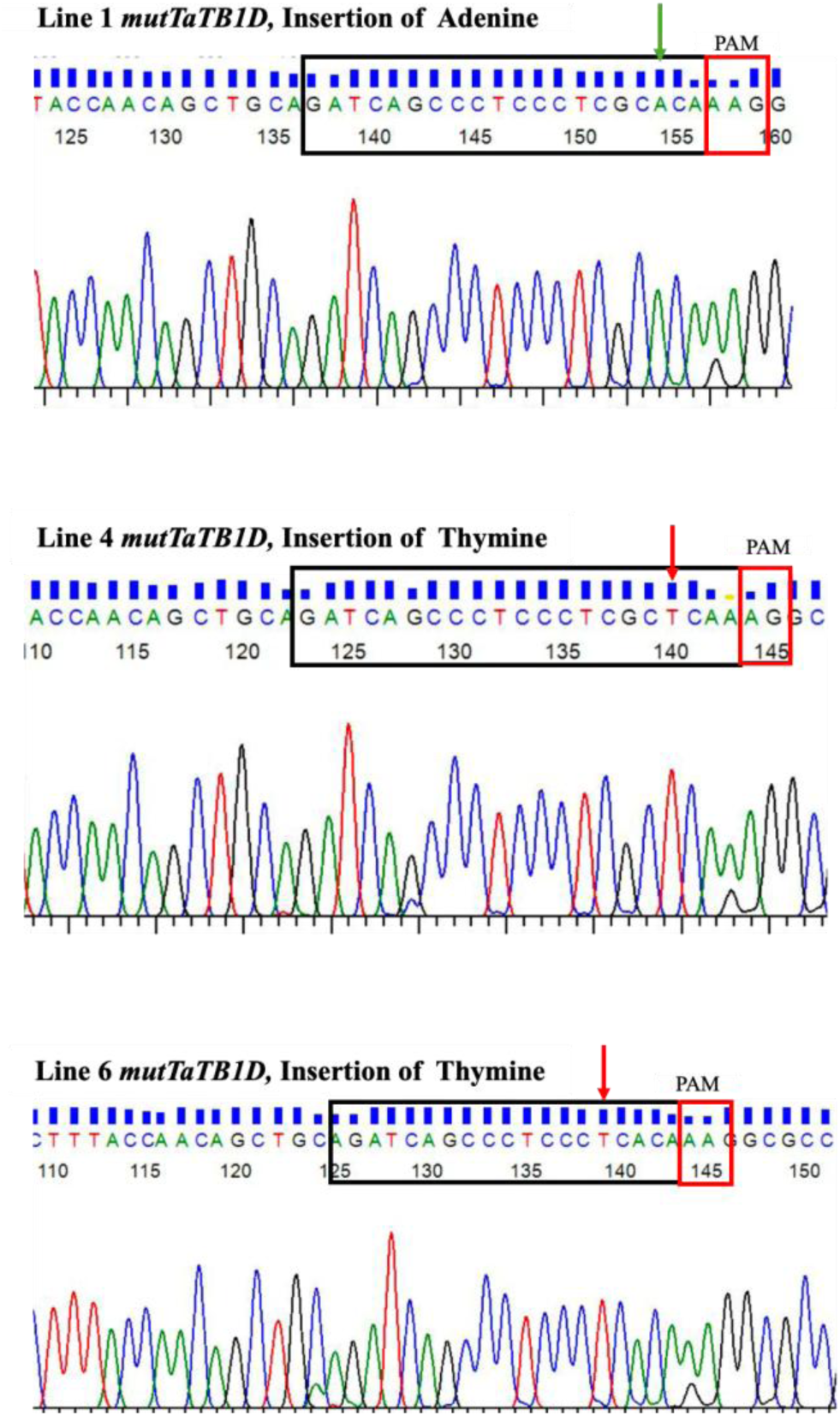
Chromatogram showing the insertion of nucleotide base adenine in mutant line 1 and thymine in mutant lines 4 and 6 of *TaTB1* at T_0_ generation.

## References

1. Awan MJA, Rasheed A, Saeed NA, Mansoor S, (2022) Aegilops tauschii presents a genetic roadmap for hexaploid wheat improvement. Trends in Genetics, 38(4), pp.307–309 10.1016/j.tig.2022.01.008

2. Cram D, Kulkarni M, Buchwaldt M, Rajagopalan N, Bhowmik P, Rozwadowski K, Parkin IA, Sharpe AG, Kagale S, (2019) WheatCRISPR: a web-based guide RNA design tool for CRISPR/Cas9-mediated genome editing in wheat. BMC plant biology, 19(1), p.474. 10.1186/s12870-019-2097-z

3. Debernardi JM, Tricoli DM, Ercoli MF, Hayta S, Ronald P, Palatnik JF, Dubcovsky J (2020) A GRF–GIF chimeric protein improves the regeneration efficiency of transgenic plants. Nature biotechnology, 38(11), pp.1274–1279. 10.1038/s41587-020-0703-0

4. Tyagi D, Banoo H, Jha DK, Meena V, Joon R, Agrwal K, Yadav P, Kumar A, Satbhai SB, Long T, Pandey AK (2026) CRISPR/Cas9 Editing of the Wheat Iron Sensor TaHRZ1 Confirms Its Conserved Role in Iron Homeostasis and Allocation in Grains. Plant, Cell & Environment 0: 1–17. 10.1111/pce.70516.

5. Dixon L E, Pasquariello M, Boden S A (2020) TEOSINTE BRANCHED1 regulates height and stem internode length in bread wheat. Journal of Experimental Botany, 71(16), 4742–4750. 10.1093/jxb/eraa252

6. Dixon LE, Greenwood JR, Bencivenga S, Zhang P, Cockram J, Mellers G, Ramm K, Cavanagh C, Swain SM, Boden SA, (2018) TEOSINTE BRANCHED1 regulates inflorescence architecture and development in bread wheat (Triticum aestivum). The Plant Cell, 30(3), pp.563–581. 10.1105/tpc.17.00961

7. Doebley J, Stec A, Gustus C, (1995) teosinte branched1 and the origin of maize: evidence for epistasis and the evolution of dominance. Genetics, 141(1), pp.333–346. 10.1093/genetics/141.1.333

8. Dong Z, Xiao Y, Govindarajulu R, Feil R, Siddoway ML, Nielsen T, Lunn JE, Hawkins J, Whipple C, Chuck G (2019) The regulatory landscape of a core maize domestication module controlling bud dormancy and growth repression. Nature communications, 10(1), p.3810.

9. Doust A (2007) Architectural evolution and its implications for domestication in grasses. Annals of Botany, 100(5), pp.941–950. 10.1038/s41467-019-11774-w

10. Dun EA, Brewer PB, Gillam EM, Beveridge CA (2023) Strigolactones and shoot branching: what is the real hormone and how does it work?. Plant and Cell Physiology, 64(9), pp.967–983. 10.1093/pcp/pcad088

11. Dun EA, de Saint Germain A, Rameau C, Beveridge CA (2012) Antagonistic action of strigolactone and cytokinin in bud outgrowth control. Plant physiology, 158(1), pp.487–498. 10.1104/pp.111.186783

12. Food and Agriculture Organization of the United Nations, 2017. The future of food and agriculture: Trends and challenges. FAO.

13. Ishizaki T, Ueda Y, Takai T, Maruyama K, Tsujimoto Y (2023) In-frame mutation in rice TEOSINTE BRANCHED1 (OsTB1) improves productivity under phosphorus deficiency. Plant Science, 330, p.111627. 10.1016/j.plantsci.2023.111627

14. Jiao Y, Wang Y, Xue D, Wang J, Yan M, Liu G, Dong G, Zeng D, Lu Z, Zhu X, Qian Q (2010) Regulation of OsSPL14 by OsmiR156 defines ideal plant architecture in rice. Nature genetics, 42(6), pp.541–544. 10.1038/ng.591

15. Jinek M, Chylinski K, Fonfara I, Hauer M, Doudna JA, Charpentier E (2012) A programmable dual-RNA–guided DNA endonuclease in adaptive bacterial immunity. Science, 337(6096), pp.816–821. 10.1126/science.1225829

16. Kerr S C, Beveridge C A (2017) IPA1: a direct target of SL signaling. Cell Res 27(10):1191–1192. 10.1038/cr.2017.114

17. Lewis JM, Mackintosh CA, Shin S, Gilding E, Kravchenko S, Baldridge G, Zeyen R, Muehlbauer GJ (2008) Overexpression of the maize Teosinte Branched1 gene in wheat suppresses tiller development. Plant Cell Reports, 27(7), pp.1217–1225. 10.1007/s00299-008-0543-8

18. Li X, Qian Q, Fu Z, Wang Y, Xiong G, Zeng D, Wang X, Liu X, Teng S, Hiroshi F, Yuan M, (2003) Control of tillering in rice. Nature, 422(6932), pp.618–621. 10.1038/nature01518

19. Lin H, Wang R, Qian Q, Yan M, Meng X, Fu Z, Yan C, Jiang B, Su Z, Li J, Wang Y (2009) DWARF27, an iron-containing protein required for the biosynthesis of strigolactones, regulates rice tiller bud outgrowth. The Plant Cell, 21(5), pp.1512–1525. 10.1105/tpc.109.065987

20. Lin Z, Chen L, Tang S, Zhao M, Li T, You J, You C, Li B, Zhao Q, Zhang D, Wang J (2023) Efficient CRISPR/Cas9-mediated genome editing in sheepgrass (Leymus chinensis). Journal of Integrative Plant Biology, 65(11), pp.2416–2420.

21. Lu Z, Yu H, Xiong G, Wang J, Jiao Y, Liu G, Jing Y, Meng X, Hu X, Qian Q, Li J (2013) Genome-wide binding analysis of the transcription activator ideal plant architecture1 reveals a complex network regulating rice plant architecture. Plant Cell 25:3743 3759. 10.1105/tpc.113.113639

22. Mohapatra PK, Sarkar RK, Panda D, Kariali E (2025) Physiology of tiller production and development. In Tillering behavior of rice plant (pp. 221–264). Singapore: Springer Nature Singapore. 10.1007/978-981-97-5235-5_8

23. Peng J, Richards DE, Hartley NM, Murphy GP, Devos KM, Flintham JE, Beales J, Fish LJ, Worland AJ, Pelica F, Sudhakar D (1999) ‘Green revolution’genes encode mutant gibberellin response modulators. nature, 400(6741), pp.256–261. 10.1038/22307

24. Ray DK, Mueller ND, West PC and Foley JA (2013) Yield trends are insufficient to double global crop production by 2050. PloS one, 8(6), p.e66428. 10.1371/journal.pone.0066428

25. Shiferaw B, Smale M, Braun HJ, Duveiller E, Reynolds M, Muricho G (2013) Crops that feed the world 10. Past successes and future challenges to the role played by wheat in global food security. Food security, 5(3), pp.291–317. 10.1007/s12571-013-0263-y

26. Song X, Lu Z, Yu H, Shao G, Xiong J, Meng X, Jing Y, Liu G, Xiong G, Duan J, Li J (2017) IPA1 functions as a downstream transcription factor repressed by D53 in strigolactone signaling in rice. Cell Res 27:1128–1141. 10.1038/cr.2017.102

27. Takeda T, Suwa Y, Suzuki M, Kitano H, Ueguchi-Tanaka M, Ashikari M, Matsuoka M, Ueguchi C (2003) The OsTB1 gene negatively regulates lateral branching in rice. The Plant Journal, 33(3), pp.513–520. 10.1046/j.1365-313X.2003.01648.x

28. Wang B, Smith SM, Li J (2018) Genetic regulation of shoot architecture. Annual review of plant biology, 69, pp.437–468. 10.1146/annurev-arplant-042817-040422

29. Wang Y, Li J (2011) Branching in rice. Current opinion in plant biology, 14(1), pp.94–99. 10.1016/j.pbi.2010.11.002

30. Zhang H, Zhang J, Wei P, Zhang B, Gou F, Feng Z, Mao Y, Yang L, Zhang H, Xu N, Zhu JK (2014) The CRISPR/C as9 system produces specific and homozygous targeted gene editing in rice in one generation. Plant biotechnology journal, 12(6), pp.797–807. 10.1111/pbi.12200

31. Zhou L, Zhang J, Yan J, Song R (2011) Two transposable element insertions are causative mutations for the major domestication gene teosinte branched 1 in modern maize. Cell research, 21(8), pp.1267–1270. 10.1038/cr.2011.104

